# Sanger validation of WGS variants - when to?

**DOI:** 10.1101/2024.04.19.590340

**Authors:** Arina Kopernik, Gaukhar Zobkova, Natalia Doroschuk, Anna Smirnova, Daria Molodtsova-Zolotukhina, Olesya Sagaydak, Oxana Ryzhkova, Sergey Kutsev, Olga Groznova, Lyusya Melikyan, Elizaveta Bondarchuk, Mary Woroncow, Eugene Albert, Viktor Bogdanov, Pavel Volchkov

## Abstract

With the development of Next-Generation Sequencing (NGS) technologies it became possible to simultaneously analyze millions of variants. Despite the quality improvement it is generally still required to confirm the variants before reporting. However, in recent years the dominant idea is that one could define the quality thresholds for “high quality” variants which do not require orthogonal validation. Despite that, no works to date report the concordance between variants from whole genome sequencing and their gold-standard Sanger validation. In this study we analyzed the concordance for 1756 WGS variants in order to establish the appropriate thresholds for high-quality variants filtering. Resulting thresholds allowed us to drastically reduce the number of variants which require validation, to 5,6% and 1.2% of the initial set for caller-agnostic thresholds and caller-dependent QUAL threshold respectively.

## Introduction

Next Generation Sequencing (NGS) is a powerful method of genome analysis. Since the introduction of the technique, according to the American College of Medical Genetics (ACMG) guidelines, discovered variants were required to be validated with an orthogonal method before reporting^1^. Usually it was done by Sanger sequencing. As NGS technologies have matured, and the volumes of sequenced samples increased, the question of the need of such validation arose. Several studies have reported small panels and exome data results with thousands of variants analyzed^4,5^,6,7, which in general showed high (91.29% - 98.7%) accordance between Sanger and NGS, reaching 100% for “high quality” variants. Hence, in recent publications^2,3^ the recommendations were somewhat relaxed: each laboratory should either establish a confirmatory testing policy for the variants or continue working with orthogonal confirmation.

Based on the abovementioned studies, the thresholds for the variant parameters were suggested as 300-500 quality score (QUAL) and 40x-100x coverage depth (DP). The most recent study^8^ concerning short variants reported that high quality variants may be considered with FILTER=PASS, QUAL >= 100, DP >= 20X and allele frequency (AF) >= 0.2%^9,10^,11. However, these recommendations, derived from the panels and exomes, seem to be of limited use for the whole genome sequencing, as its mean depth of coverage is usually around 30x.

During our research, we could not encounter studies reporting validation results on the whole genome data, and the minority of all the reported data considers BGI technological platform. The aim of this study was to perform the analysis of the quality control parameters for *1756* potentially causative variants derived from WGS data of 1150 patients. Each of these variants was validated by Sanger sequencing. Based on that analysis we suggest a set of quality filters capable of separating variants which require Sanger validation from the “high quality” variants (the ones which do not require validation) with F1-score of 99,5%. Implementing these metrics into routine practice will greatly reduce the need for Sanger sequencing, therefore decreasing time and cost of overall clinical WGS analysis.

## Results

### Dataset

A cohort of 1150 WGS samples was analyzed to find the cause of the disease and to identify carriers of potentially pathogenic variants. In total 1756 variants were subjected to further validation, out of them 1555 SNVs and 201 INDELS, 1555 variants located in exons, 201 in introns. Mean coverage of the samples was 34.1, ranging from 20.57 to 48.64. Mean coverage depth (DP) of the variants was 33 (3-81), quality (QUAL) was 492 (30-2106)

All selected 1756 variants were subjected to validation by Sanger sequencing. In 5 (0, 28%) cases WGS data did not match Sanger data, demonstrating 99.72% concordance.

### Optimal thresholds selection

Quality parameters were compared with the thresholds in recent publications^4,5^,6,7 (300-500 quality score and 40x-100x depth coverage) in order to demonstrate whether they are applicable to WGS data. In this work we will analyze thresholds from the most recent manuscript^8^. Thresholds presented there (FILTER = PASS, QUAL > 100, DP > 20, AF > 0.2) allowed to filter out 250 “low-quality” variants, 5 of which were unconfirmed resulting in 2% specificity.

It is possible to divide presented thresholds into two groups: caller-agnostic (AF and DP) and caller-specific (QUAL) parameters based on whether they depend on variant calling results or not. Speaking of caller-agnostic thresholds, variants with allele frequency (AF) parameter > 0.2 and DP > 20 alone demonstrated 100% concordance with Sanger data while “low-quality” variants showed 2% specificity detecting unconfirmed variants. It is possible to lower DP parameter to 15 and increase AF to 0.25 in order to increase specificity from 2% (250 low-quality variants, 5 of them unconfirmed) to 5% (99 low-quality variants, 5 of them unconfirmed) without losing sensitivity.

As for quality parameter (QUAL), it can be analyzed separately as a caller-specific threshold. All the variants above 100 demonstrated 100% concordance with Sanger data while 5 out of 21 variants with QUAL lower than 100 were unconfirmed which resulted in 76% concordance with NGS. (Figure 1).

**Figure 1.**
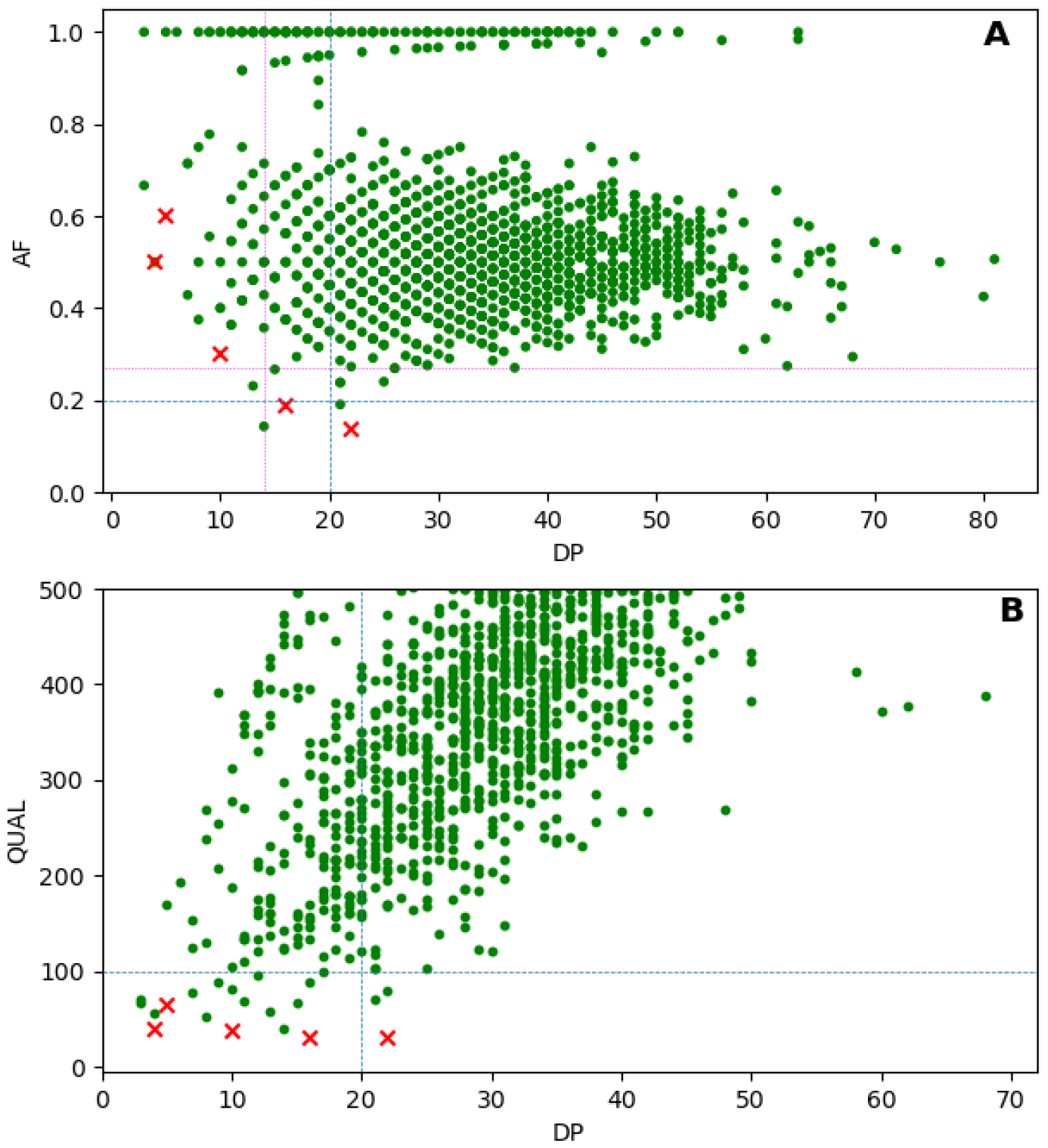
(A) Distribution of confirmed and unconfirmed variants depending on AF and DP parameters. (B) Distribution of confirmed and unconfirmed variants depending on QUAL and DP parameters. Green points represent confirmed variants and red X-s represent unconfirmed variants. Blue lines represent thresholds from recent WES study, magenta line represents caller-agnostic quality thresholds suggested in this work (DP > 15, AF > 0.25)

The results presented in Table 1 represent the ratio of high-quality to low-quality variants both for the thresholds from the exome study^8^ and for our data. An increased specificity for thresholds of AF and DP (from 2% to 5%) is observed due to the decreased number of low-quality variants (from 249 to 99). For QUAL parameter alone there were only 21 low-quality variants, including the unconfirmed ones which resulted in 23,8% (21 low-quality variants, 5 of them unconfirmed) specificity. Additionally, the F1-scores for both sets of thresholds were calculated in order to compare them. The old set of thresholds turned out to have 0.925 while the new set of caller-independent parameters increased this metric to 0.972. F1-score for QUAL threshold resulted in 0.995.

**Table 1.** Classification statistics based on different thresholds

| Quality thresholds | “High quality” variants (unconfirmed variants) | “Low quality” variants (unconfirmed variants) | Sensitivity | Specificity | F1-score |
| --- | --- | --- | --- | --- | --- |
| FILTER = PASS<br>QUAL > 100<br>DP > 20<br>AF > 0.2<sup>8</sup> | 1506 (0) | 250 (5) | 100% | 2% | 0.925 |
| DP > 15<br>AF > 0.25 | 1657 (0) | 99(5) | 100% | 5% | 0.972 |
| QUAL > 100 | 1735 (0) | 21(5) | 100% | 23,8% | 0.995 |

## Discussion

The main aim of the validation policy for NGS data is to minimize the amount of additional testing without lowering the quality of a final result. Hence, the goal of the study was to identify a set of thresholds which robustly separate good quality WGS variants from the ones that require validation with appropriate specificity and sensitivity.

According to Table 1, previously suggested threshold values for clinical exomes^8^ work reasonably well on WGS data, filtering out all the unconfirmed variants into the “low quality” bin. However, because of the differences in average sequencing depth between WES and WGS, the threshold of DP > 20 is too high for WGS data, hence the specificity of such approach is lower than one might expect (2%), which results in F1-score of 0.925 Additionally, QUAL and FILTER parameters are caller-specific and therefore the values are not strictly comparable between bioinformatic pipelines.

Based on our data, better results can be obtained using only caller-agnostic parameters of DP and AF. DP > 15 and AF > 0.25 achieve similarly sensitive results, filtering out all 5 unconfirmed variants into the “low quality” bin while shrinking the bin itself 2.5 times, which, if implemented as an actual policy, would reduce the confirmatory testing cost accordingly.

However, an even greater result can be achieved using the QUAL parameter alone. Setting a single QUAL > 100 achieves the 23,8% specificity and F1 score of 0.995 without the reduction in sensitivity for unconfirmed variants, which shrinks the “low quality” bin size to less than 2% of the total dataset size. This result is understandable since the QUAL parameter itself encompasses a complex set of rules aimed at providing an evaluation of confidence in the presence of the variant at a given site. Of note, we would not recommend a direct transfer of this threshold to different callers apart from the one used in this work (HaplotypeCaller v.4.2)

## Conclusions

Our study demonstrates that previously suggested thresholds (DP>20, AF>0.2 QUAL>100) work reasonably well for WGS data separating the false variants with 100% sensitivity, 2% specificity and F1-score resulting in 0.925. However, for WGS we suggest lowering the DP requirements for “high quality” variants and achieve, in our case, 5% specificity with the same sensitivity and increasing F1-score to 0.972 for caller-agnostic thresholds (DP > 15, AF > 0.25). Additionally, QUAL is shown to be an independent classification parameter which alone, for our dataset, can separate false variants with much greater specificity of 23,8% and F1-score 0.995 using QUAL=100 threshold for HaplotypeCaller V4.2 thus tremendously reducing the need for validation.

## Materials and methods

### Samples and Library preparation

Library preparation was performed using PCR-free enzymatic shearing protocol (MGIEasy FS PCR-Free DNA Library Prep Kit, MGI, China). Whole genome sequencing of 1150 patients in PE150 mode was performed using DNBseq-T7 and DNBseq-G400 (MGI, China) sequencers at EVOGEN LLC laboratory (Moscow, Russia).All the experimental stages were carried out in accordance with the manufacturer’s standard protocols. The average whole genome sequencing depth was 30×. The variant calling of the genetic variants was carried out using bioinformatics analysis accelerators (EVA Pro (EVOGEN, Russia) and MegaBOLT (MGI, China)).

### Bioinformatics pipeline

Raw reads were trimmed with fastp (0.23.1) mapped to the reference human genome, build hg38 with minimap2 (2.17). Calling was performed after base recalibration with haplotype caller (4.2) in a single sample mode according to GATK best practice guidelines.

### Variant selection

After the bioinformatic analysis the data were reviewed by clinical variant scientists and geneticists in the search for the disease-causing variants according to ACMG Standards and Guidelines, considered variants were pathogenic, likely pathogenic and uncertain significance. The candidate variants for the reporting were assigned to the lab for Sanger sequencing validation.

### Sanger Sequencing

Causative genetics variants revealed by WGS were validated by Sanger sequencing at EVOGEN LLC. Sample DNA extraction was performed using QIAamp DNA blood Mini Kit (QIAGEN, Germany) / MGIEasy Magnetic Beads (MGI, China) by standard manufacturer’s protocols. Purified PCR products were sequenced using the BigDye Terminator Kit v3.1 and ABI 3500 Genetic Analyzer (Applied Biosystems, United States) by standard manufacturer’s protocols. The analysis of the sequenced data was performed using the software Variant Reporter Software v3.0 (Applied Biosystems, United States).

### Data analysis

Information on quality, allele frequency and depth coverage were extracted from vcf files. The dataset was analyzed with the custom python scripts. For the evaluation of the thresholds the F1-score was used as a metric that balances precision and recall. Precision is calculated as a number of true positive variants (high quality and confirmed with Sanger) divided by the sum of true positive and false positive (high quality and unconfirmed with Sanger) variants. Recall is calculated as the number of true positive variants divided by the sum of true positive and false negative (low quality and confirmed with Sanger) variants. The specificity and sensitivity are calculated according to the following formulas:

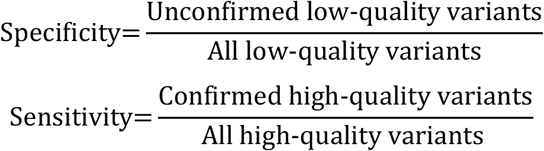

## Data availability

The data that support the findings of this study are available from Evogen LLC but restrictions apply to the availability of these data, which were used under license for the current study, and so are not publicly available. Data are however available from the authors upon reasonable request and with permission of Evogen LLC.

## Acknowledgements

Whole genome sequencing, data analysis for selected samples and variant validation by Sanger sequencing were performed in the Evogen LLC by laboratory staff (Antonenko A., Aydarova V., Belov R., Bartsis A., Boldireva B., Bykadorov P., Dibirova H., Domoratskaya E., Frolkov A, Hahanova V., Gubona M., Golovanova M., Leonova V., Markov D., Mikhaylov V., Nikitina O., Panferova A., Pudova L., Revkova M., Rodionova D., Safina S., Shichkova A., Sokolova N., Ulanova P., Vasilenko A., Veselova G., Zolotopup A., deputy lab head Krinitsina A., and lab head Belenikin M.).

The research was supported by the Ministry of Science and Higher Education of the Russian Federation (agreement # *№* 075-03-2022-107/10)

## Author information

Affiliations

^1^Federal Research Center for Innovator and Emerging Biomedical and Pharmaceutical Technologies, 125315 Moscow, Russia

^2^Evogen LLC, Moscow, Russia

^3^Federal State Budgetary Institution “Research Centre For Medical Genetics”, 115478 Moscow, Russia.

^4^Lomonosov Moscow State University, Moscow, Russia

^5^Veltischev Research and Clinical Institute for Pediatrics and Pediatric Surgery on the Pirogov Russian National Research Medical University of the Ministry of Health of the Russian Federation

^6^Charity Fund for medical and social genetic aid projects «Life Genome»

^7^The Pirogov Russian National Research Medical University of the Ministry of Health of the Russian Federation

Corresponding author

Correspondence to Viktor Bogdanov and Pavel Volchkov.

## Contributions

P.Y., V.B. conceived the experiment. G.Z., N.D., A.S., D.Z., O.S. reviewed the bioinformatic analysis. A.K., V.B., and E.A. calculated the statistical data and devised the thresholds. All authors participated in the discussion and reviewed the manuscript.

## Additional information

### Competing interests

The authors declare no competing interests

## Notes

### Competing Interest Statement

The authors have declared no competing interest.

